## Extended Data for "Cryo-EM Structure of a Mammalian-specific Alternative Amyloid Exon"

SUPPLEMENTARY TABLES

**Supplementary Table 1.** Cryo-EM data collection and refinement statistics.

| **Name** | **hnRNPDL-2 amyloid fibril** |
|---|---|
| **PDB ID**<br>**EMDB ID**<br>**EMPIAR** | **7ZIR**<br>**EMDB-14738**<br>**EMPIAR-11064** |
| **Data collection** | |
| Magnification | ×120000 |
| Pixel size (Å) | 0.889 |
| Defocus range (μm) | -1.0 to -2.5 |
| Voltage (kV) | 200 |
| Camera | Falcon 3EC |
| Microscope | Talos Arctica |
| Number of frames | 40 |
| Total dose ($e^-$/Å$^2$) | 40 |
| **Reconstruction** | |
| Micrographs | 1114 |
| Manually picked fibrils | 10450 |
| Box size (pixel) | 380 |
| Inter-box distance (Å) | 33.25 |
| Segments extracted | 158493 |
| Segments after Class2D | 158493 |
| Segments after Class3D | 54490 |
| Resolution (Å) | 2.5 |
| Map sharpening B-factor (Å$^2$) | -46.0 |
| Helical rise (Å) | 4.82 |
| Helical twist (°) | - 4.86 |
| **Atomic model** | |
| Non-hydrogen atoms | 2120 |
| Protein residues | 255 |
| Waters | 50 |
| r.m.s.d. Bond lengths | 0.001 (0) |
| r.m.s.d. Bond angles | 0.300 (0) |
| All-atom clash score | 2.48 |
| Rotamers outliers | 0 |
| Ramachandran outliers | 0 |
| Ramachandran allowed | 2.04 |
| Ramachandran favored | 97.96 |

**Supplementary Table 2.** Solvent-accessible surface area (SASA) parameters calculated for the upper layers of hnRNPDL-2, hnRNPA2, hnRNPA1, hSAA1, Aβ-42 and $\alpha$-synuclein amyloid fibrils.

| | Total Upper Surface (Å$^2$) | Buried Upper Surface (Å$^2$) | Exposed Upper Surface (Å$^2$) | Exposed Polar Surface (%) |
|---|---|---|---|---|
| **hnRNPDL-2** | 5789 | 2330 | 3458 | 48 |
| **hnRNPA1** | 4721 | 2480 | 2241 | 42 |
| **hnRNPA2** | 5761 | 2382 | 3379 | 42 |
| **hSAA1** | 6482 | 3078 | 3404 | 38 |
| **$\alpha$-synuclein** | 6168 | 2521 | 3648 | 36 |
| **Aβ-42** | 3644 | 1706 | 1938 | 32 |

Note that proteins are ordered according to decreasing Exposed Polar Surface (%) values. Exposed Polar (%) values were calculated using GetArea[1]. Surfaces values were calculated with PDBePISA[2]. PDB accession codes used for calculations are: hnRNPDL-2 (PDB 7ZIR), hnRNPA2 (PDB 6WQK), hnRNPA1 (PDB 7BX7), hSAA1 (PDB 6MST), Aβ-42 (PDB 5KK3) and $\alpha$-synuclein (PDB 6OSJ).

**Supplementary Table 3.** Solvent-accessible surface area (SASA) parameters calculated for the inner layers of hnRNPDL-2, hnRNPA2, hnRNPA1, hSAA1, Aβ-42 and $\alpha$-synuclein amyloid fibrils.

| | Total Inner Surface (Å$^2$) | Buried Inner Surface (Å$^2$) | Exposed Inner Surface (Å$^2$) | Exposed Polar Surface (%) |
|---|---|---|---|---|
| **hnRNPDL-2** | 5761 | 4467 | 1294 | 57 |
| **hnRNPA2** | 5758 | 4518 | 1240 | 47 |
| **hnRNPA1** | 4721 | 4117 | 604 | 44 |
| **hSAA1** | 6477 | 5474 | 1003 | 43 |
| **Aβ-42** | 3589 | 2868 | 720 | 40 |
| **$\alpha$-synuclein** | 6170 | 4741 | 1429 | 37 |

Note that proteins are ordered according to decreasing Exposed Polar Surface (%) values. Exposed Polar (%) values were calculated using GetArea[1]. Surfaces values were calculated with PDBePISA[2]. PDB accession codes used for calculations are: hnRNPDL-2 (PDB 7ZIR), hnRNPA2 (PDB 6WQK), hnRNPA1 (PDB 7BX7), hSAA1 (PDB 6MST), Aβ-42 (PDB 5KK3) and $\alpha$-synuclein (PDB 6OSJ).

**Supplementary Table 4.** Solvation free energy gain ($\Delta G^{int}$) parameters calculated for hnRNPDL-2, hnRNPA2, hnRNPA1, $\alpha$-synuclein, hSAA1 and Aβ-42 amyloid fibrils.

| | **$\Delta G^{int}$ (kcal/mol)** | **$\Delta G^{int}$/residue (kcal/mol)** |
|---|---|---|
| **hnRNPDL-2** | -13 | -0,251 |
| **hnRNPA2** | -15 | -0,257 |
| **hnRNPA1** | -13 | -0,292 |
| **$\alpha$-synuclein** | -20 | -0,325 |
| **hSAA1** | -18 | -0,338 |
| **Aβ-42** | -16 | -0,512 |

Note that proteins are ordered according to increasing $\Delta G^{int}$/residue values. ΔG values were calculated with PDBePISA[2]. PDB accession codes used for calculations are: hnRNPDL-2 (PDB 7ZIR), hnRNPA2 (PDB 6WQK), hnRNPA1 (PDB 7BX7), $\alpha$-synuclein (PDB 6OSJ), hSAA1 (PDB 6MST) and Aβ-42 (PDB 5KK3).

**Supplementary Table 5.** Calculated Gibbs free energy values upon mutations on hnRNPDL-2, hnRNPA1 and hnRNPA2 amyloid fibrils.

| **Protein** | **Mutation** | **ΔΔG (kcal/mol)** |
|---|---|---|
| **hnRNPDL-2** | D259H | 2.50 |
| | D259N | -5.54 |
| **hnRNPA1** | D262N | -15.60 |
| | D262V | -29.84 |
| **hnRNPA2** | D290V | -9.33 |

### EXTENDED DATA FIGURES

#### Extended Data Fig. 1

```
hnRNPDL-1   1- MEVPPRLSHVPPPLFPSAPATLASRSLSHWRPRPPRQLAPLLPSLAPSSARQGARRAQRH -60
hnRNPDL-2      ------------------------------------------------------------
hnRNPDL-3      ------------------------------------------------------------

hnRNPDL-1  61- VTAQQPSRLAGGAAIKGGRRRRPDLFRRHFKSSSIQRSAAAAAATRTARQHPPADSSVTM -120
hnRNPDL-2   1- -----------------------------------------------------------M -1
hnRNPDL-3   1- -----------------------------------------------------------M -1

hnRNPDL-1 121- EDMNEYSNIEEFAEGSKINASKNQQDDGKMFIGGLSWDTSKKDLTEYLSRFGEVVDCTIK -180
hnRNPDL-2   2- EDMNEYSNIEEFAEGSKINASKNQQDDGKMFIGGLSWDTSKKDLTEYLSRFGEVVDCTIK -61
hnRNPDL-3   2- EDMNEYSNIEEFAEGSKINASKNQQDDGKMFIGGLSWDTSKKDLTEYLSRFGEVVDCTIK -61

hnRNPDL-1 181- TDPVTGRSRGFGFVLFKDAASVDKVLELKEHKLDGKLIDPKRAKALKGKEPPKKVFVGGL -240
hnRNPDL-2  62- TDPVTGRSRGFGFVLFKDAASVDKVLELKEHKLDGKLIDPKRAKALKGKEPPKKVFVGGL -121
hnRNPDL-3  62- TDPVTGRSRGFGFVLFKDAASVDKVLELKEHKLDGKLIDPKRAKALKGKEPPKKVFVGGL -121

hnRNPDL-1 241- SPDTSEEQIKEYFGAFGEIENIELPMDTKTNERRGFCFITYTDEEPVKKLLESRYHQIGS -300
hnRNPDL-2 122- SPDTSEEQIKEYFGAFGEIENIELPMDTKTNERRGFCFITYTDEEPVKKLLESRYHQIGS -181
hnRNPDL-3 122- SPDTSEEQIKEYFGAFGEIENIELPMDTKTNERRGFCFITYTDEEPVKKLLESRYHQIGS -181

hnRNPDL-1 301- GKCEIKVAQPKEVYRQQQQQQKGGRGAAAGGRGGTRGRGRGQGQNWNQGFNNYYDQGYGN -360
hnRNPDL-2 182- GKCEIKVAQPKEVYRQQQQQQKGGRGAAAGGRGGTRGRGRGQGQNWNQGFNNYYDQGYGN -241
hnRNPDL-3 182- GKCEIKVAQPKEVYRQQQQQQKGGRGAAAGGRGGTRGRGRGQ------------------ -223
                                                                 NLS
hnRNPDL-1 361- YNSAYGGDQNYSGYGGYDYTGYNYGNYGYGQGYADYSGQQSTYGKASRGGGNHQNNYQPY -420
hnRNPDL-2 242- YNSAYGGDQNYSGYGGYDYTGYNYGNYGYGQGYADYSGQQSTYGKASRGGGNHQNNYQPY -301
hnRNPDL-3      ---------------------------------------QSTYGKASRGGGNHQNNYQPY -244
```

**Extended Data Fig. 1. Sequence alignment of hnRNPDL isoforms.** The amino acid sequences of hnRNPDL from three different majoritarian isoforms (isoform 1, hnRNPDL-1; isoform 2, hnRNPDL-2 and isoform 3, hnRNPDL-3) were aligned with Clustal Omega. Uniprot accession numbers are: hnRNPDL-1 (O14979-1), hnRNPDL-2 (O14979-2) and hnRNPDL-3 (O14979-3). The residues comprising the N-terminal Arg-rich LCD and RRMs are shown in orange and blue, respectively. The amino acids comprising the Gly/Tyr-rich amyloid exon 6 are depicted in green. The Asp residue in position 259 (D259), which is mutated in LGMDD3 patients, is shown in red. The C-terminal amino acids that encode for the hnRNPDL nuclear localization signal (NLS) are indicated.

**Extended Data Fig. 2**

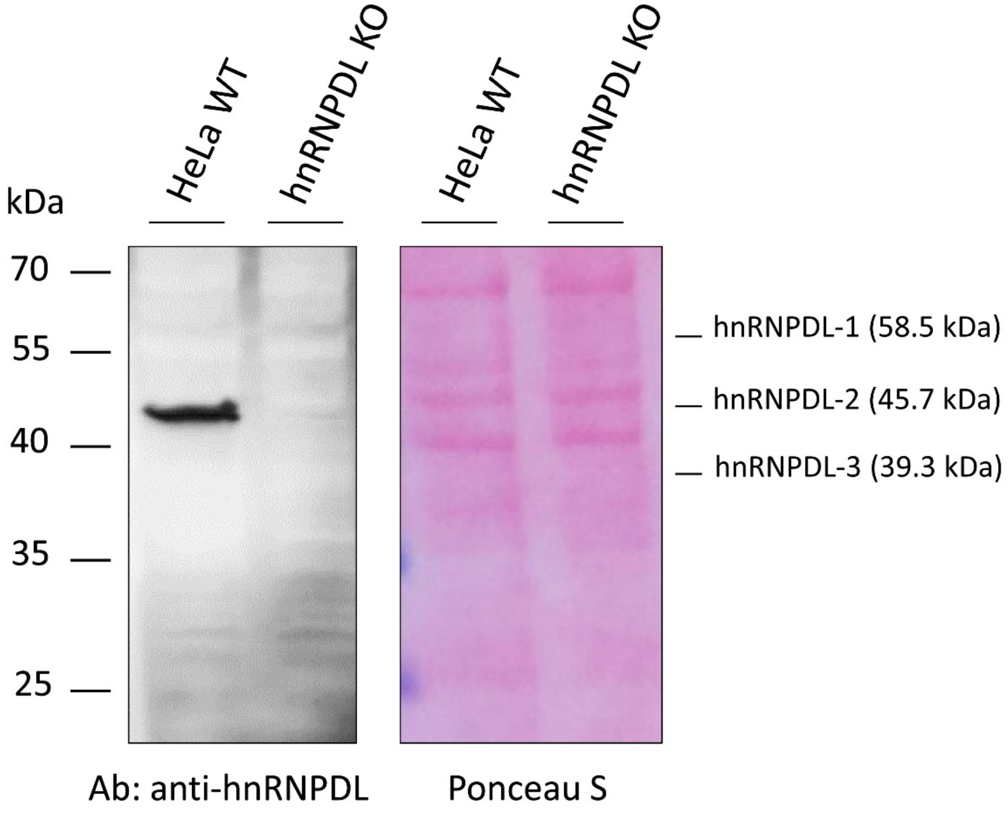

**Extended Data Fig. 2. Characterization of HeLa hnRNPDL knockout cell line.** Western blot analysis of the hnRNPDL expression levels in wild-type (WT) and hnRNPDL KO HeLa cells (left panel). Ponceau S stain is shown as loading control (right panel).

#### Extended Data Fig. 3

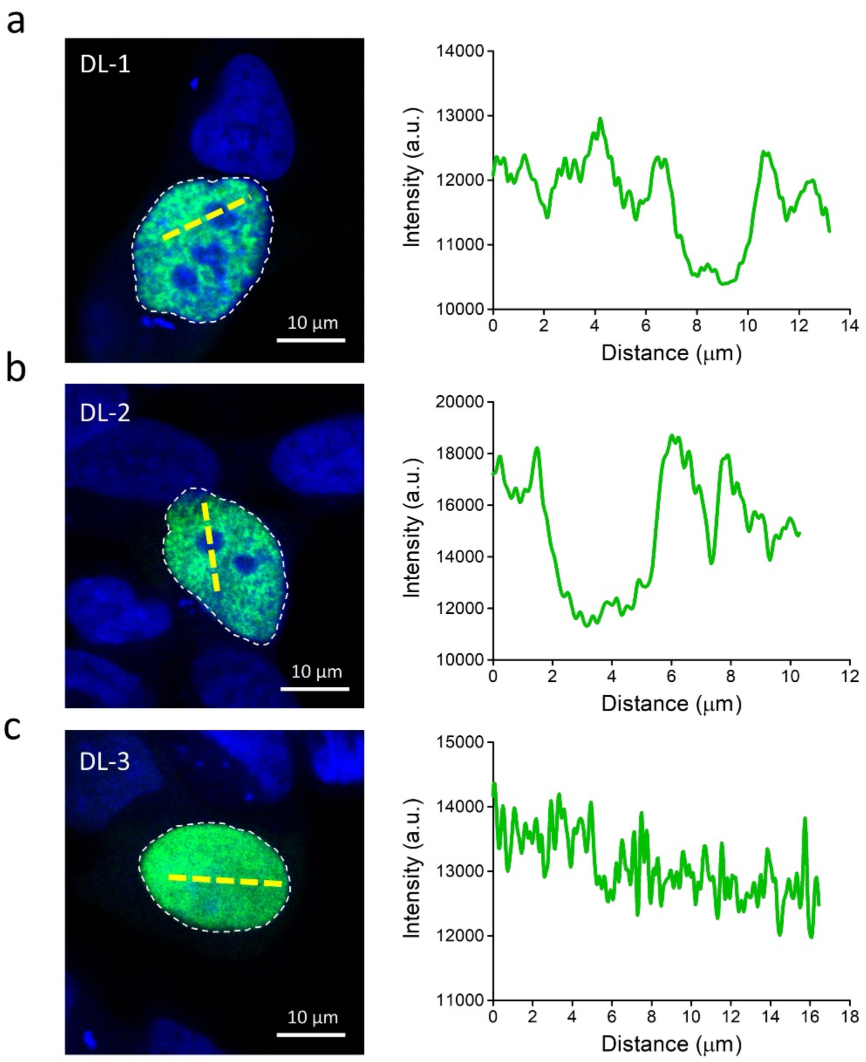

**Extended Data Fig. 3. Nuclear localization of GFP-tagged hnRNPDL isoforms in HeLa cells.** Confocal laser scanning microscopy (CLSM) images (left panels) and intensity profile analyses (right panels) of HeLa cells transiently transfected with the three different hnRNPDL isoforms (GFP-hnRNPDL-1 (a), GFP-hnRNPDL-2 (b) and GFP-hnRNPDL-3 (c)).

#### Extended Data Fig. 4

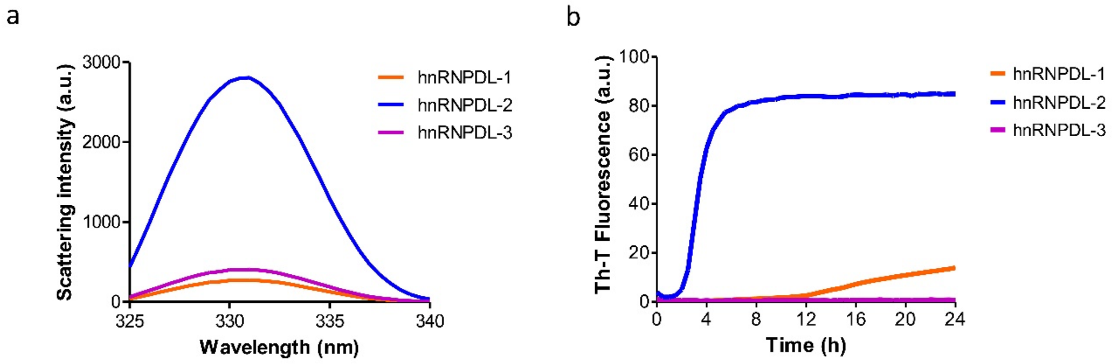

**Extended Data Fig. 4. hnRNPDL isoforms amyloid formation.** a) Light scattering analysis of hnRNPDL-1 (orange line), hnRNPDL-2 (blue line) and hnRNPDL-3 (purple line) aggregates after incubation at 37 °C under shaking at 600 rpm for 48 h. b) Aggregation kinetics of hnRNPDL-1 (orange line), hnRNPDL-2 (blue line) and hnRNPDL-3 (purple line) at 25 µM. Aggregation is shown as Th-T binding over time and samples were incubated at 37 °C and under constant shaking in 96 well plates. In b), data is expressed as mean ± s.e.m (n = 3 independent experiments).

#### Extended Data Fig. 5

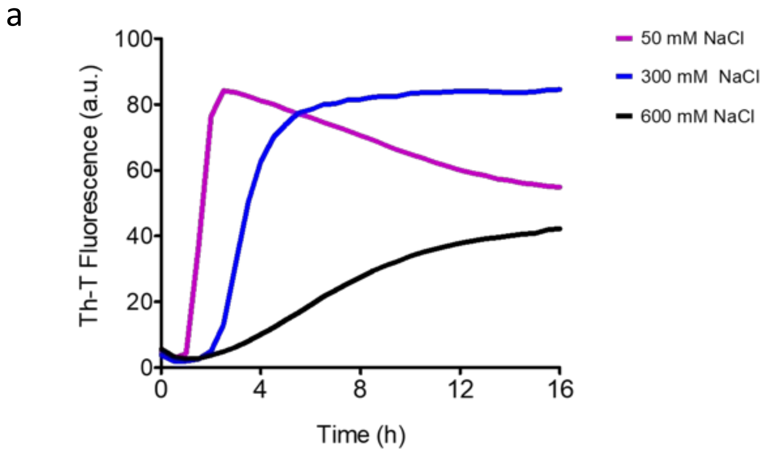

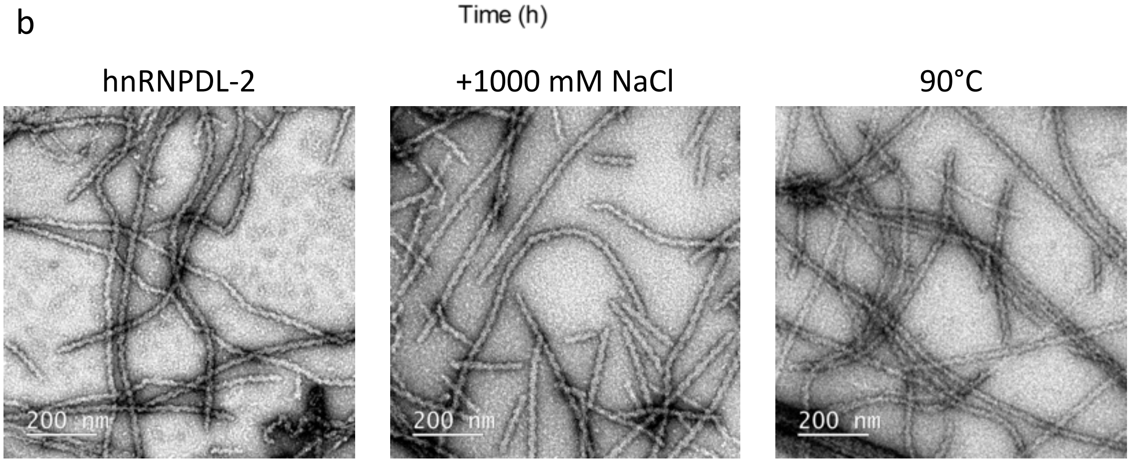

**Extended Data Fig. 5. Effect of NaCl and temperature on hnRNPDL-2 aggregation and fibril stability.** a) Aggregation kinetics of soluble hnRNPDL-2 at three different NaCl concentrations. Aggregation is shown as Th-T binding over time. Samples were incubated at 37 °C and under constant shaking in 96 well plates as described. b) Representative negative-staining TEM micrographs of pre-formed hnRNPDL-2 amyloid fibrils after 48 h in the absence (left) or presence (middle) of 1000 mM NaCl, and after 2h at 90 °C (right).

**Extended Data Fig. 6**

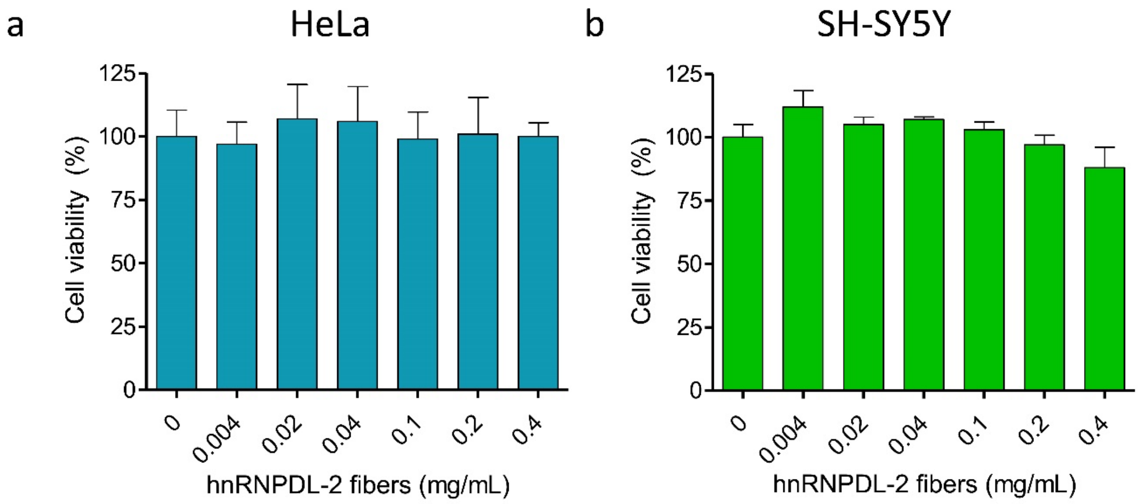

**Extended Data Fig. 6. hnRNPDL-2 forms non-toxic amyloid fibrils.** a, b) Effect of hnRNPDL-2 amyloid fibrils on the viability of HeLa (a) or human neuroblastoma SH-SY5Y (b) cells after incubation with the pre-formed filaments for 48 h.

**Extended Data Fig. 7**

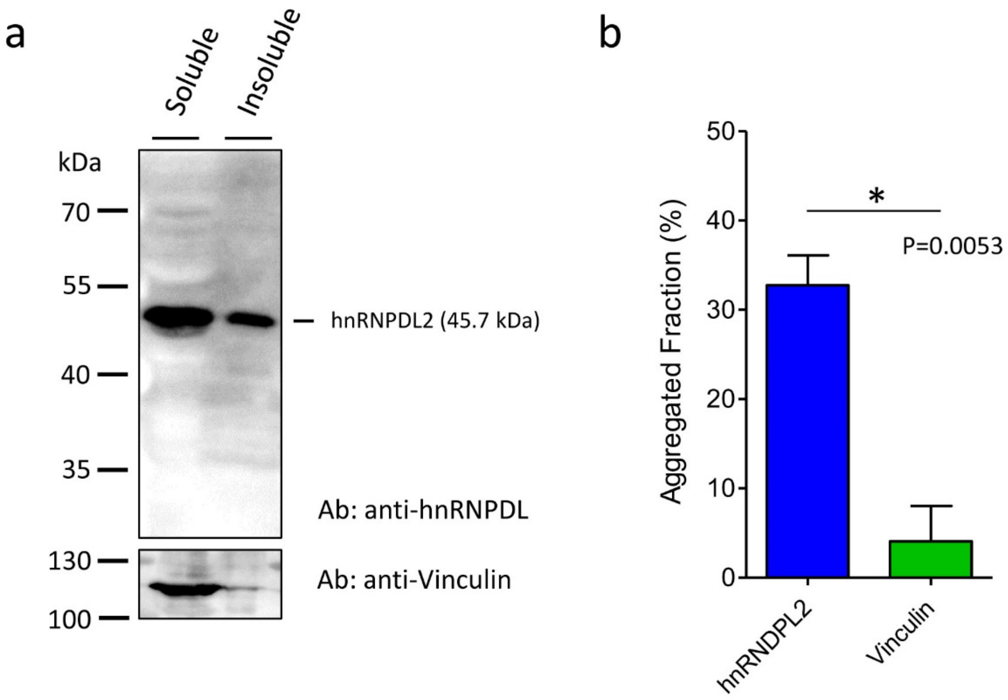

**Extended Data Fig. 7. Detergent-soluble and insoluble fractions of cellular endogenous hnRNPDL-2 in HeLa cells.** a) Western blot analysis of the soluble and insoluble fraction of HeLa cells lysed with mPER Mammalian Protein Extraction Reagent. The presence of hnRNPDL-2 in soluble and insoluble fractions was detected by immunoblotting with anti-hnRNPDL antibody. The presence of vinculin in the different fractions was determined as control protein. b) Relative Levels (%) of hnRNPDL-2 or vinculin present in the aggregated fraction (detergent-insoluble fraction) of HeLa cells after cell lysis.

**Extended Data Fig. 8**

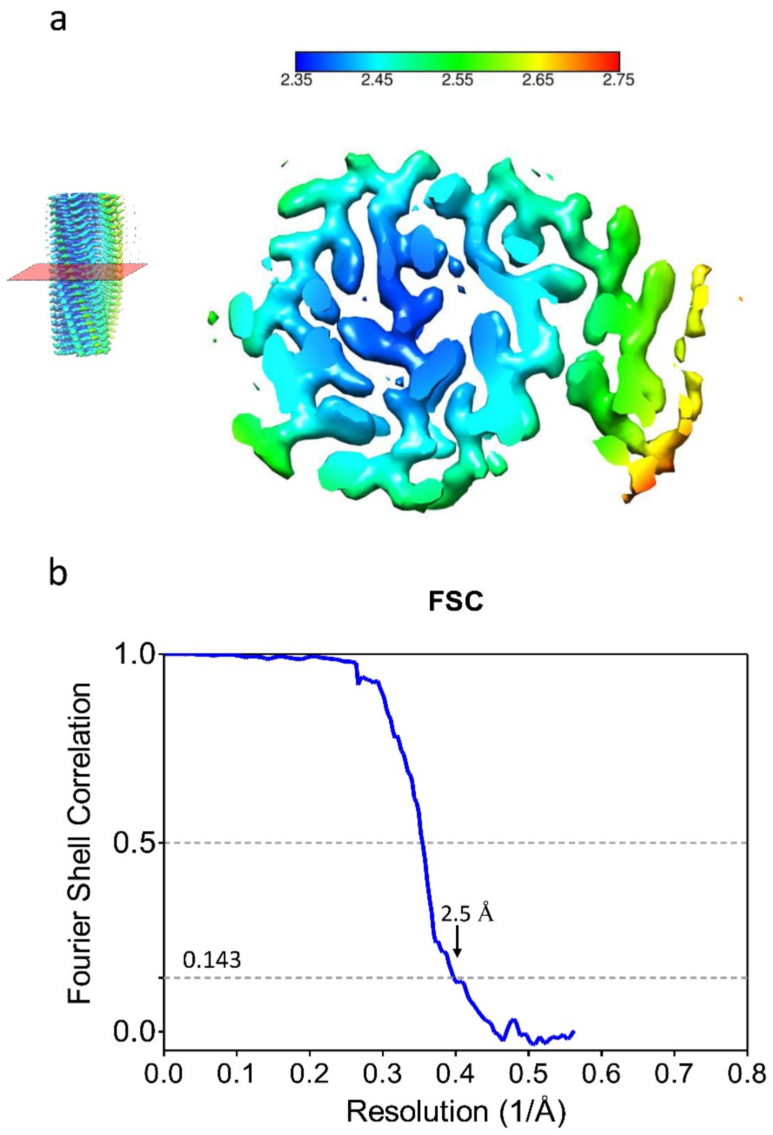

**Extended Data Fig. 8. Resolution estimation of hnRNPDL-2 amyloid filaments.** a) Local resolution map of the hnRNPDL-2 filament reconstruction; local resolution levels (Å) are color coded on top. b) Fourier Shell Correlation curve (FSC) of hnRNPDL-2 amyloid filaments. The overall resolution of the best reconstruction is 2.5 Å (FSC=0.143).

**Extended Data Fig. 9**

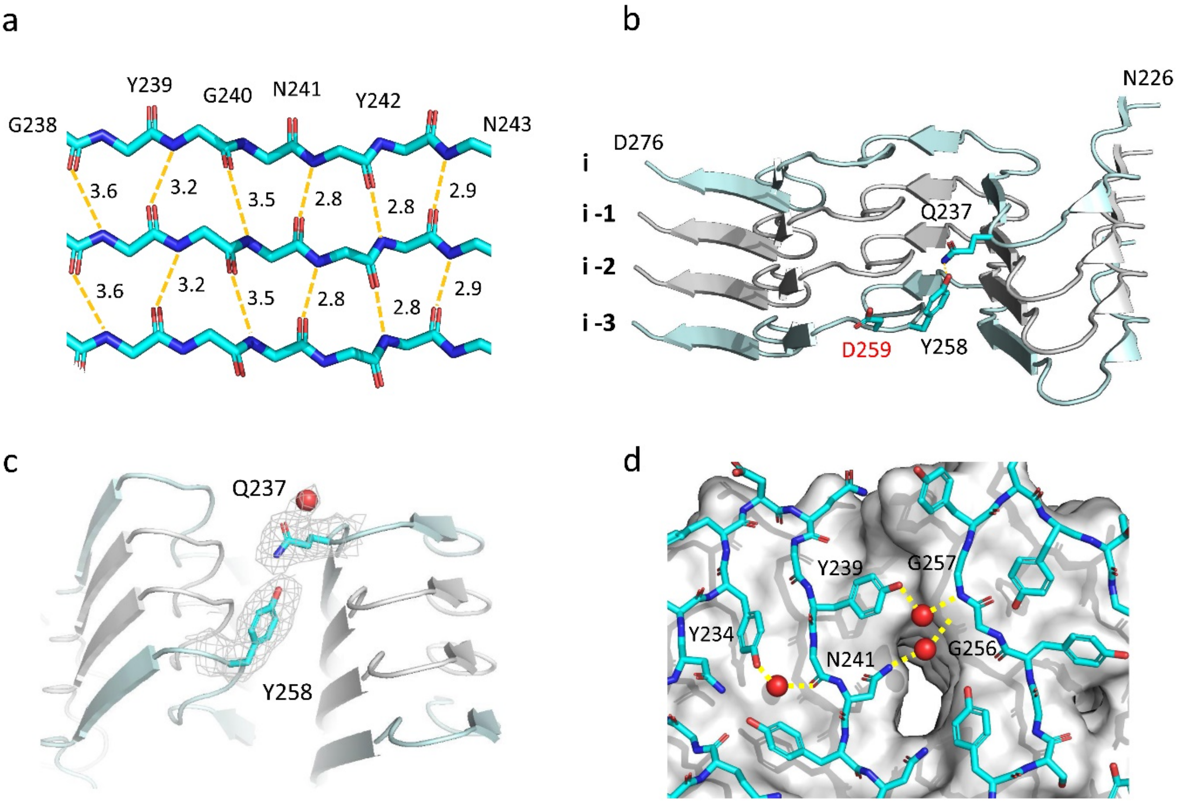

**Extended Data Fig. 9. Relevant interactions in the hnRNPDL-2 protofilament interface.** a) Close-up view of the inter-molecular hydrogen-bonding network holding consecutive parallel β-strands between residues G238 and N243. b) Side view of four layers of the amyloid core of hnRNPDL-2, depicting the interaction between the side chains of Q237 and Y258. The side chain of D259, which is mutated in LGMDD3, is also shown as sticks. c) Close up view of the residues involved in the interaction between the layers i and i -3. The side chains of Q237 and Y258 are shown as sticks colored in cyan over the cryo-EM map sown as a grey mesh. d) Close-up view of the central water channel found in the hnRNPDL-2 amyloid core. Note that the hydrogen bonds between the two water molecules in the channel of the first layer and the adjacent residues are depicted with yellow dashed lines.

#### Extended Data Fig. 10

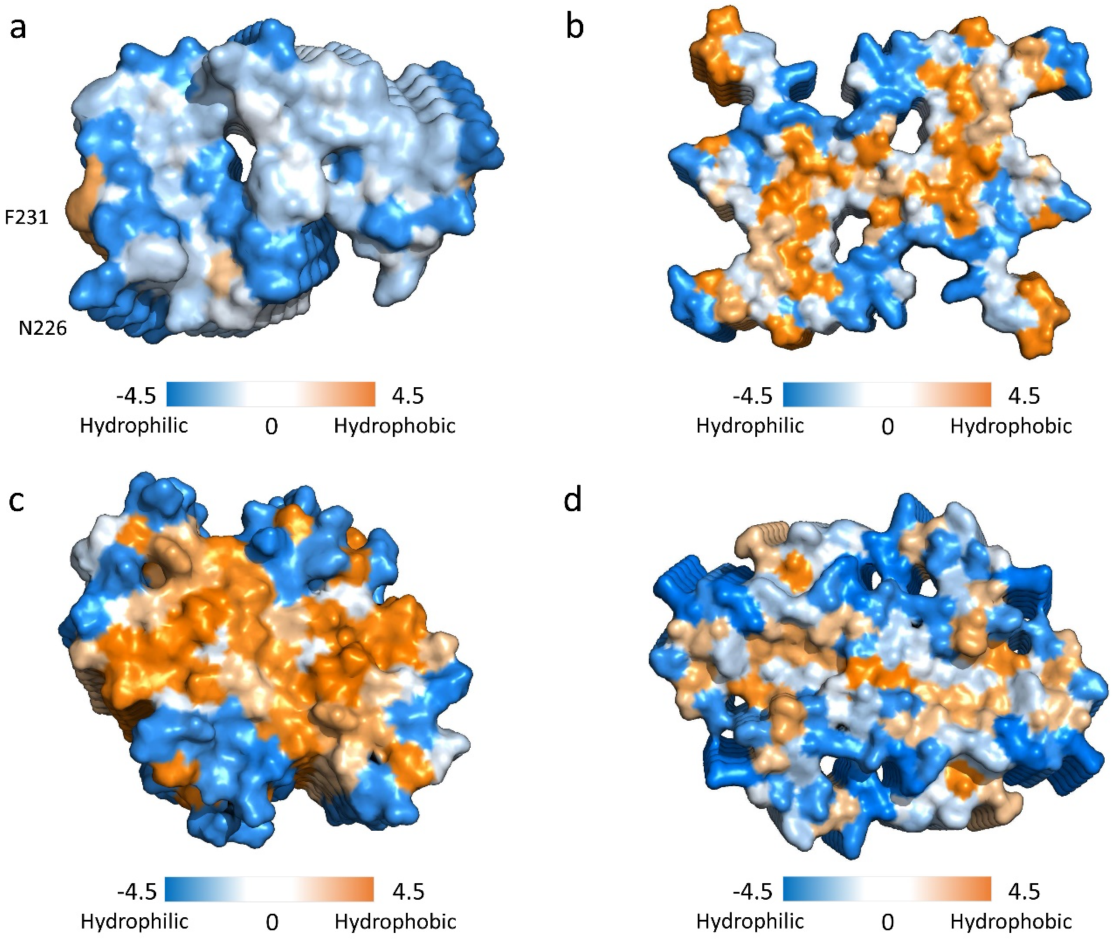

**Extended Data Fig. 10. hnRNPDL-2 fibrils display a highly polar surface interface.** Surface representation of a) hnRNPDL-2 (PDB 7ZIR), b) α-synuclein (PDB 6OSJ), c) Amyloidβ (Aβ)-42 (PDB 5KK3) and d) human serum amyloid A (PDB 6MST) fibrils cross-sections colored with the hydrophobicity levels of each residue. The hydrophobicity levels were assigned to each residue according to the Kyte-Doolittle scale[3]. Note that a unique strong hydrophobic spot is visible in the core of hnRNDPL-2, which corresponds to the side chain of F231 near the N-terminus.

#### Extended Data Fig. 11

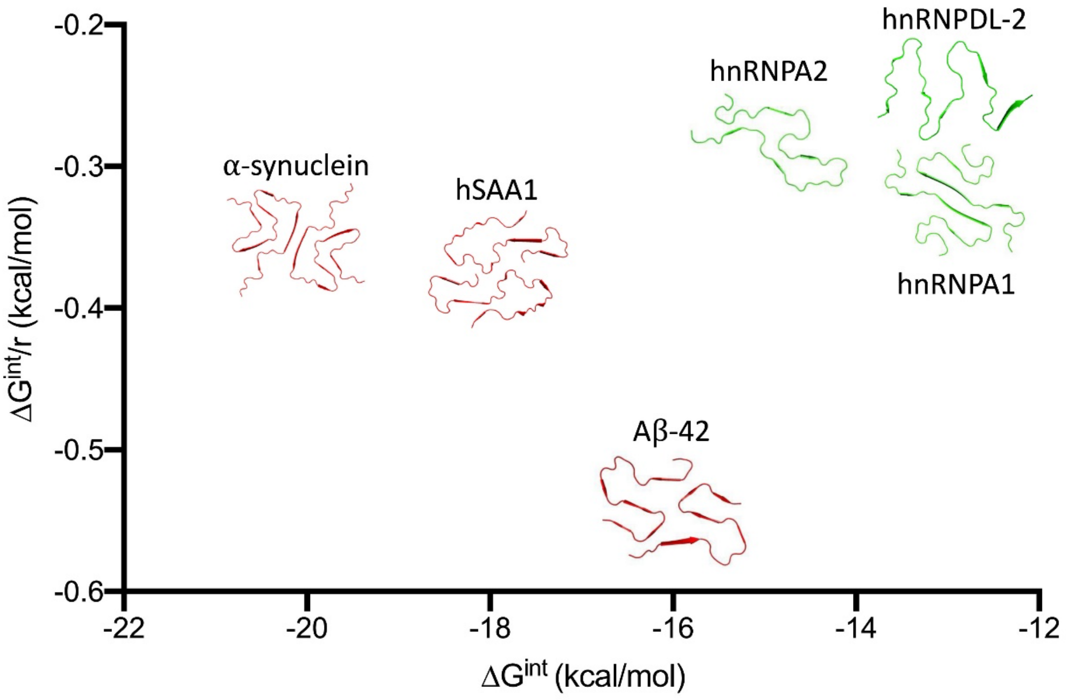

**Extended Data Fig. 11. Stability of hnRNPDL-2, hnRNPA2, hnRNPA1, *α*-synuclein, hSAA1 and Aβ-42 amyloid fibrils.** Represented with fibril structures in two dimensions: solvation free energy gain per molecule (horizontal) and per residue (vertical). Amyloid fibril structures that are non-pathogenic and less stable are colored in green, whereas pathogenic and more stable ones are colored in red. PDB codes are: hnRNPDL-2 (PDB 7ZIR), hnRNPA2 (PDB 6WQK), hnRNPA1 (PDB 7BX7), *α*-synuclein (PDB 6OSJ), hSAA1 (PDB 6MST) and Aβ-42 (PDB 5KK3).

#### Extended Data Fig. 12

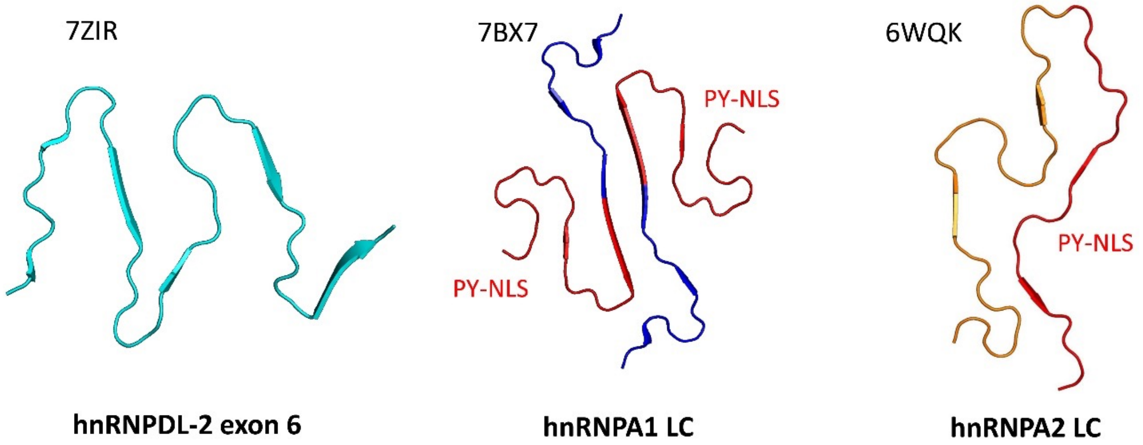

**Extended Data Fig. 12. Comparison of cryo-EM structures of amyloid fibrils from RNA/DNA binding proteins.** Ribbon representation of the cryo-EM amyloid core structures formed by the low complexity regions of hnRNPDL-2, hnRNPA1 and hnRNPA2. In hnRNPA1/A2, the regions corresponding to the nuclear localization sequence (PY-NLS) are colored in red. PDB codes are indicated over each structure.

#### Extended Data Fig. 13

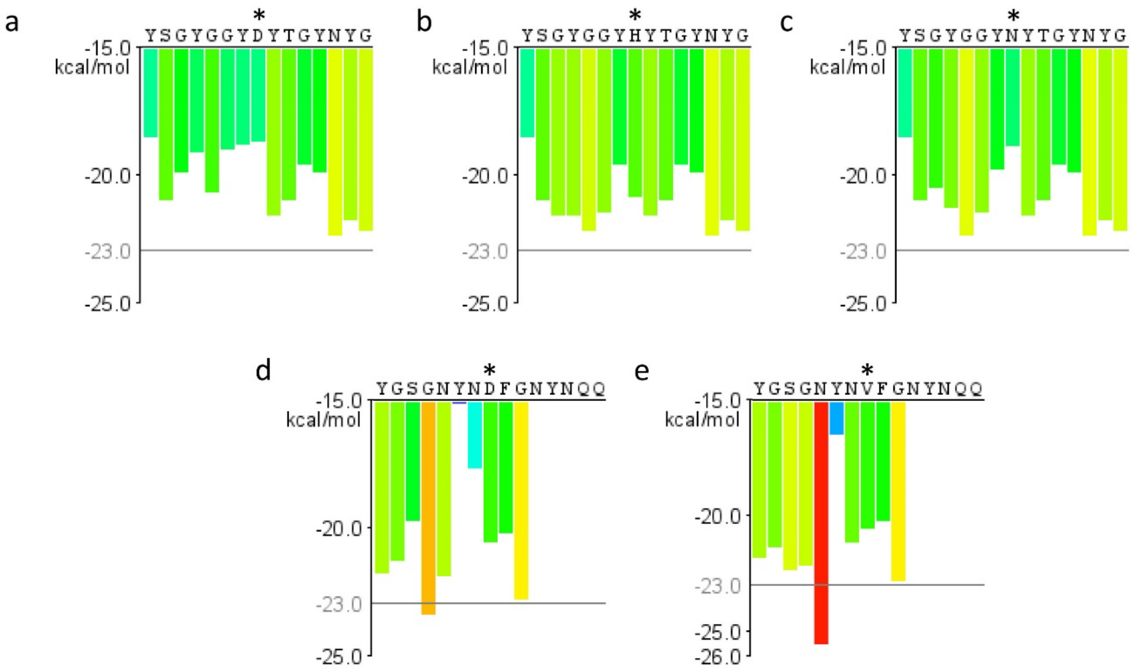

**Extended Data Fig. 13. Predicted energy for fibrillation of six-residue windows of hnRNPDL-2 and hnRNPA2 disease-associated mutants.** a) hnRNPDL-2 WT, b) hnRNPDL-2 D259H, c) hnRNPDL-2 D259N, d) hnRNPA2 WT and e) hnRNPA2 D290V. Residues which are mutated in disease are indicated with an asterisk. Red bars represent hexapeptides with energy below -23kcal/mol, predicted to form fibrils. Data obtained from ZipperDB[4].

#### Extended Data Fig. 14

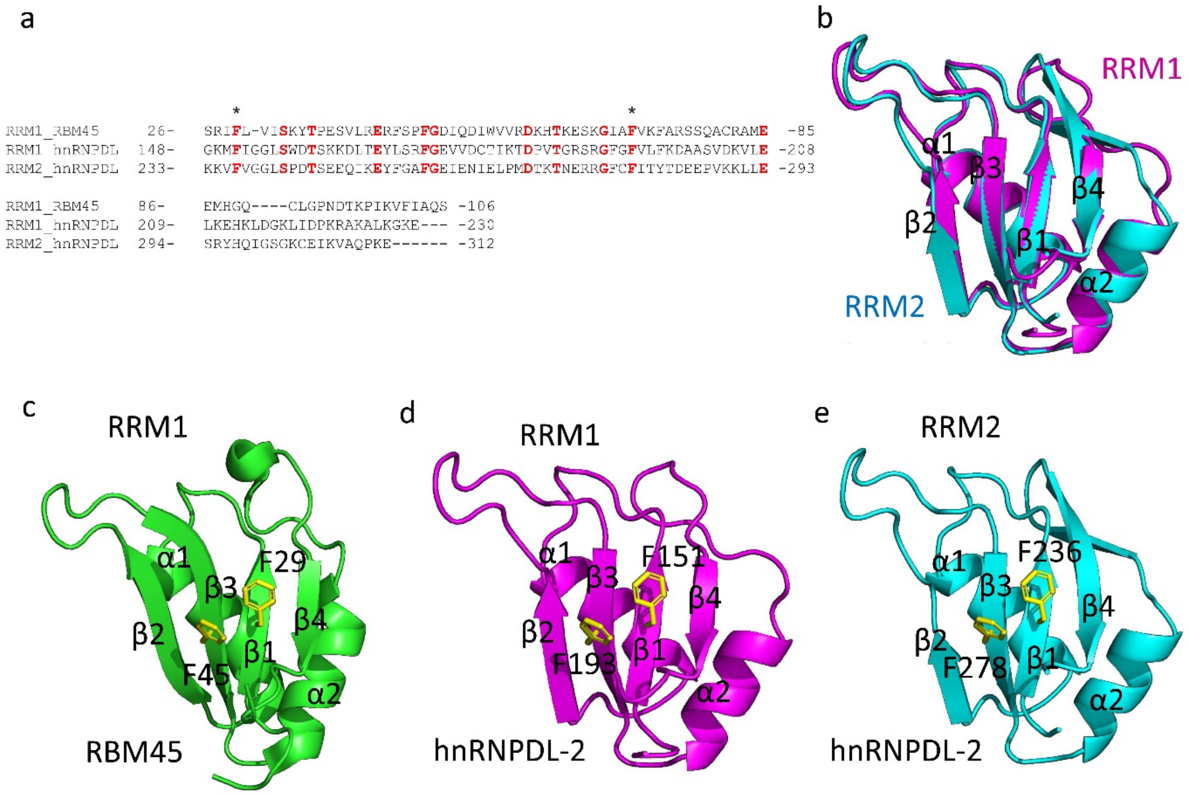

**Extended Data Fig. 14. Structure of the RNA/DNA-binding domains from hnRNPDL-2.** a) Sequence alignment of both hnRNPDL-2 RRMs with the RRM1 from RBM45. Identical residues are highlighted in red. The positions of aromatic residues that participate in RNA/DNA binding are indicated with an asterisk. b) Structural alignment of RRM1 and RRM2 from hnRNPDL-2. The structural models were generated with AlphaFold2[5]. c-e) Structural comparison of the first RRM domain from RBM45 (PDB 7CSX) (c) and both RRM1 (d) and RRM2 (e) from hnRNPDL-2. In c-e) the side chains of two of the conserved aromatic residues found in the RNA/DNA binding pocket are shown in yellow sticks.

#### Extended Data Fig. 15

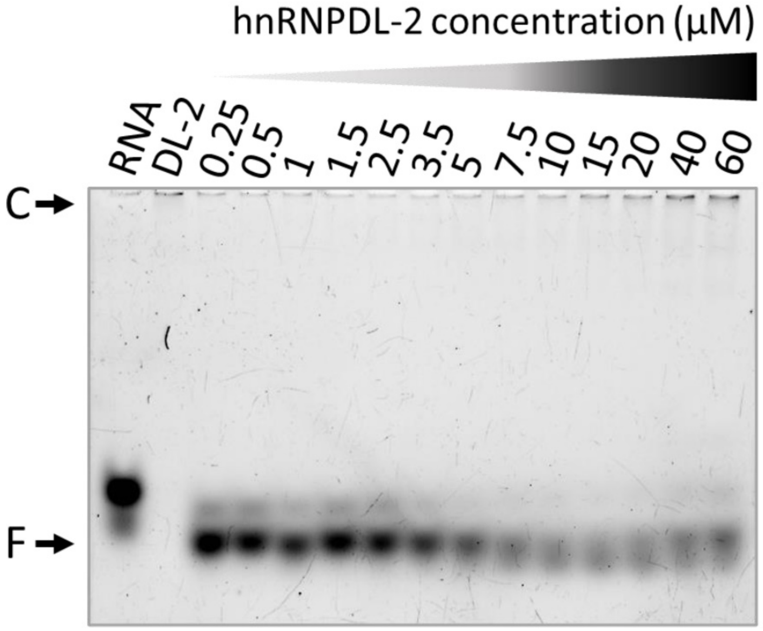

**Extended Data Fig. 15. RNA (F-GACUAGC) binding to soluble hnRNPDL-2.** Electrophoretic mobility shift assay (EMSA) of soluble hnRNPDL-2 with a Fluorescein-labelled RNA (F-GACUAGC). The 7-mer RNA was incubated with increasing concentrations of the soluble form of hnRNPDL-2. The assayed protein concentrations are indicated in µM.

#### Extended Data Fig. 16

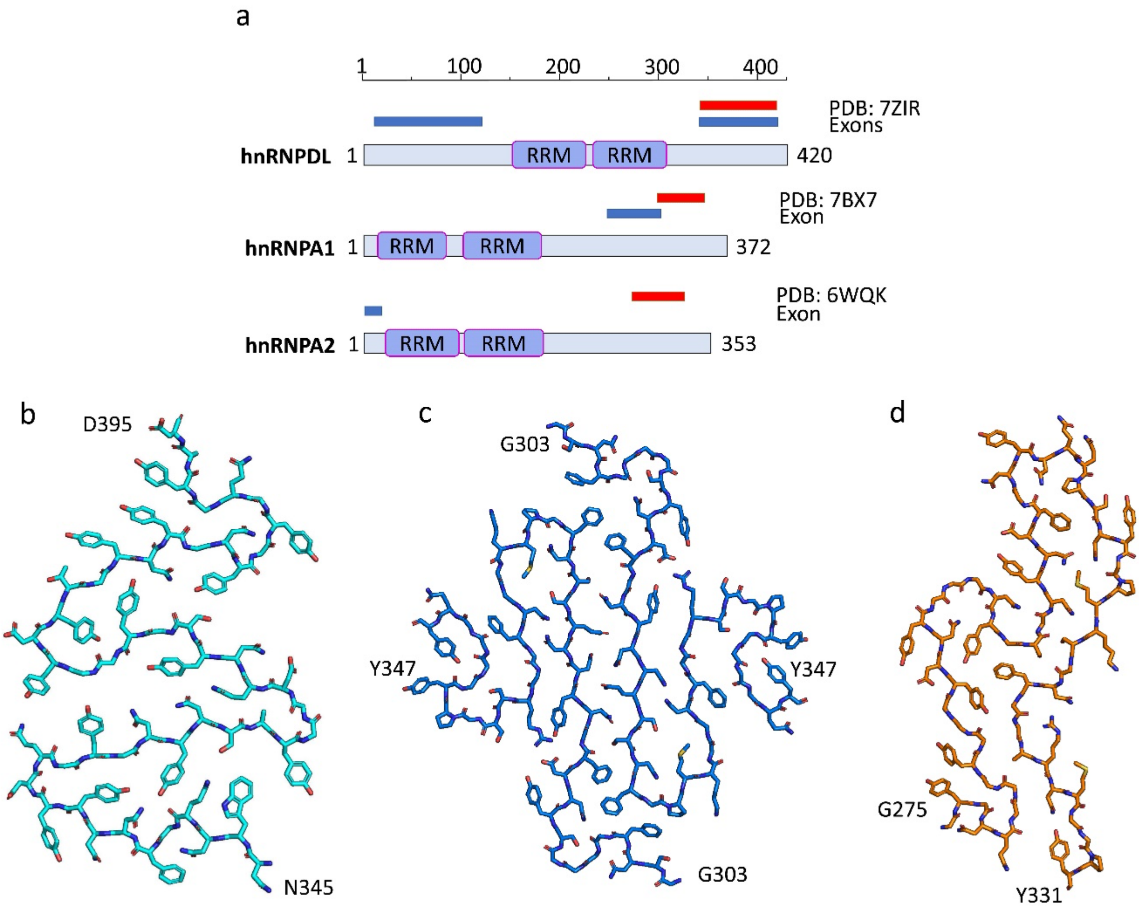

**Extended Data Fig. 16. Location of amyloid cores and conserved exons in hnRNPDL, hnRNPA1 and hnRNPA2.** a) Primary structures of hnRNPDL, hnRNPA1 and hnRNPA2 indicating the location of the solved amyloid cores on top (red boxes). The position of exonic regions (exon/s) are also indicated over the protein structures (blue boxes). These regions correspond to the major splicing isoforms described in the Uniprot Database[6]. b-d) Atomic models of the amyloid cores of b) hnRNPDL-2 (PDB 7ZIR), c) hnRNPA1 (PDB 7BX7) and d) hnRNPA2 (PDB 6WQK).

#### EXTENDED DATA REFERENCES
